# Distinct Transcriptomic Responses to Aβ plaques, Neurofibrillary Tangles, and *APOE* in Alzheimer’s Disease

**DOI:** 10.1101/2023.03.20.533303

**Authors:** Sudeshna Das, Zhaozhi Li, Astrid Wachter, Srinija Alla, Ayush Noori, Aicha Abdourahman, Joseph A. Tamm, Maya E. Woodbury, Robert V. Talanian, Knut Biber, Eric H. Karran, Bradley T. Hyman, Alberto Serrano-Pozo

## Abstract

**INTRODUCTION:** Omics studies have revealed that various brain cell types undergo profound molecular changes in Alzheimer’s disease (AD) but the spatial relationships with plaques and tangles and *APOE*-linked differences remain unclear.

**METHODS:** We performed laser capture microdissection of Aβ plaques, the 50μm halo around them, tangles with the 50μm halo around them, and areas distant (>50μm) from plaques and tangles in the temporal cortex of AD and control donors, followed by RNA-sequencing.

**RESULTS:** Aβ plaques exhibited upregulated microglial (neuroinflammation/phagocytosis) and downregulated neuronal (neurotransmission/energy metabolism) genes, whereas tangles had mostly downregulated neuronal genes. Aβ plaques had more differentially expressed genes than tangles. We identified a gradient Aβ plaque>peri-plaque>tangle>distant for these changes. AD *APOE*ε4 homozygotes had greater changes than *APOE*ε3 across locations, especially within Aβ plaques.

**DISCUSSION:** Transcriptomic changes in AD consist primarily of neuroinflammation and neuronal dysfunction, are spatially associated mainly with Aβ plaques, and are exacerbated by the *APOE*ε4 allele.

## INTRODUCTION

Alzheimer’s disease (AD) is increasingly viewed as a complex neurodegenerative disorder in which not only neurons but also glial and vascular cells undergo profound morphological, molecular, and functional changes, potentially due to chronic exposure to its two defining pathological hallmarks: amyloid-β (Aβ) plaques and phospho-tau (pTau) neurofibrillary tangles (NFTs)^1, 2^. For example, reactive astrocytes and microglia are spatially associated with Aβ plaques and NFTs^3–7^, and dense-core Aβ plaques are frequently decorated by pTau-positive dystrophic neurites^7^ and associated with synaptic loss^8^. Furthermore, whether and how the apolipoprotein E (*APOE*) genotype—the main genetic modifier of AD risk—affects the microenvironment of Aβ plaques and NFT-bearing neurons and the cellular responses to these lesions are crucial questions that could help explain the substantial heterogeneity in rate of cognitive decline that characterizes AD^9–11^. A number of single-nucleus RNA-sequencing (snRNA-seq) studies on postmortem brain samples have reported transcriptomic differences of various cell types between AD and control (CTRL) donors, but snRNA-seq lacks the spatial information necessary to address the question of whether the transcriptomic alterations in neurons and glia occur primarily in close proximity to Aβ plaques and NFTs, or whether they are diffuse throughout the cortex^12–18^. Spatial transcriptomic methods have been successfully applied to study the transcriptomic changes around Aβ plaques in AD mouse models^19, 20^ but their application in human postmortem AD brains is technically challenging and has limitations^21^. Thus, a better understanding of the transcriptional changes in the vicinity of Aβ plaques and NFTs and the possible mediator role of the *APOE* genotype is critical to resolve this complexity and identify potential therapeutic targets to develop drugs capable of impacting Aβ aggregation or clearance, pTau aggregation and/or propagation, and downstream neurodegeneration.

Here we performed laser capture microdissection (LCM) of Aβ plaques and NFTs from postmortem brain cryostat sections of CTRL and AD donors followed by bulk RNA-seq to investigate the transcriptomic changes occurring within and around Aβ plaques and NFTs in the AD brain as well as the effect of the *APOE* genotype on those changes. Specifically, we tested the hypotheses that (1) transcriptomic differences between AD and CTRL donors are maximum within and around Aβ plaques and NFTs; (2) Aβ plaques and NFTs are associated with distinct transcriptomic changes in the AD brain; (3) the *APOE* genotype differentially impacts the transcriptomic changes associated with Aβ plaques and NFTs in the AD brain.

## METHODS

### Human Donors

Frozen tissue from the superior temporal gyrus (BA22) of n=8 control and n=10 AD (n=5 *APOE*ε4/ε4, n=4 *APOE*ε3/ε3, and n=1 *APOE*ε3/ε4) donors was obtained from the Massachusetts Alzheimer’s Disease Research Center (MADRC) Brain Bank. Donors were selected based on frozen tissue availability, RNA Integrity Number (RIN) ≥5 (2100 Bioanalyzer, Agilent), and in the AD group, *APOE* genotype. CTRL donors were cognitively normal before death and had Braak NFT stage 0-III and CERAD neuritic plaque (NP) score none or sparse. AD donors met current clinical diagnostic criteria of dementia due to probable AD^22^ and current neuropathological diagnostic criteria of definite AD, that is, had a CERAD NP score of moderate or frequent and a Braak NFT stage of V/VI^23, 24^. All donors were free of other potentially confounding neurological and neurodegenerative diseases as reflected by review of clinical information and complete neuropathological examination. All donors or their next-of-kin provided written informed consent for the brain donation and the present study was approved under the MGH Institutional Review Board.

### Laser capture microdissection

Fifteen-micron-thick cryostat sections were mounted onto non-polarized glass slides (Gold Seal Rite-On Ultra Frost, Thermo-Scientific), thawed, slightly fixed in 75% ethanol for 40 sec, and stained with Thioflavin-S 0.05% in 100% ethanol (ThioS, Sigma-Aldrich, Cat# 1326-12-1) for 5 min, followed by dehydration in increasing concentrations of ethanol and xylene, and air dried. Immediately, sections were placed onto the stage of an Arcturus Veritas LCM apparatus (Arcturus, Thermo-Scientific) and laser capture microdissection (LCM) was performed with a laser power of 80-85 μV and pulse duration of 3,500 μsec to fill up ten CapSure LCM Macro caps (Thermo-Scientific, Cat# LCM0211) per region of interest and per donor. In CTRL donors, approximately 100 mm^2^ of cortex were dissected. In AD tissues, four different regions of interest were dissected: ThioS+ plaques (approx. 1,000/donor), the 50 μm area around those ThioS+ plaques, ThioS+ NFTs with the 50 μm area around them (approx. 1,500/donor), and cortex far (>50 μm) from both ThioS+ plaques and NFTs (approx. 80-100 mm^2^).

### RNA extraction and purification

Caps containing laser-capture microdissected tissue were covered with 20 μL RNA extraction buffer for 30 min at 42°C and RNA extracts were spun down at 800x*g* for 2 min and stored at -80°C until further use. Extracted RNA was purified using the Arcturus PicoPure RNA Isolation kit (ThermoFisher Scientific, Cat# KIT0204), following manufacturer’s instructions. Briefly, the ten RNA extracts from each region of interest of each donor were treated with DNase and consolidated in the same mini-column. Purified RNA was eluted from the mini-columns in two steps of 7 μL. Quality of purified RNA was confirmed using the 6000 RNA Pico kit (Agilent, Cat# 5067-1513) in a 2100 Bioanalyzer instrument (Agilent, Cat# G2939BA), with RIN ranging between 2.0 and 5.2, and DV200 (i.e., the % of RNA fragments longer than 200 nucleotides) between 70 and 95%.

### Library preparation and RNA-sequencing

cDNA libraries were prepared using the SMARTer^®^ Stranded Total RNA-Seq Kit v2 – Pico Input Mammalian (Takara Bio) following manufacturer’s instructions in ten batches. Batch effects were controlled with three measures: (1) Sample allocation to plates was randomized; (2) External RNA Controls Consortium (ERCC) RNA Spike-In Mixes (Ambion, Life Technologies) consisting of 92 poly-adenylated transcripts with known concentration were added to each RNA sample; and (3) Same reference sample was loaded in all plates.

A pilot study conducted in the NextSeq550 Illumina platform with single-end sequencing (75 bp), dual index, 13 cycles, rendered an average number of input reads of 57.7M with a 40% average GC content, and high duplicate rates, therefore the number of amplification cycles was reduced. For the final study, sequencing was conducted in a NextSeq2000 Illumina platform with paired-end sequencing (2×60bps), dual index, and 11 cycles, rendering an average number of input reads of 33.9M. Sequence alignment was conducted against the Homo *sapiens* genome assembly GRCh38.

### Sequencing alignment and quality control

Sequencing quality control was performed with the FastQC (version 0.11.9)^25^ and MultiQC (version 1.9)^26^ software. Alignment was conducted with STAR (version 2.7.1a)^27^ against the Homo *sapiens* genome assembly GRCh38 (gencode v31). FeatureCounts (v1.6.5)^28^ was used to assign aligned reads to genes. Genes lowly expressed across conditions were filtered out, retaining those expressed at ≥1 CPMs in at least 4 samples. Raw counts were log-transformed and trimmed mean of M values (TMM) normalized^29^. After removal of duplicate reads expected at high rates from low input RNA samples, downstream analysis was performed based on a mean library size of 10.72M reads. Principal Component Analysis (PCA) of normalized counts demonstrated a good separation of CTRL and AD samples with a good separation by *APOE* genotype that was not driven by the CTRL group. The reference sample included in all sequencing batches and the ERCC spike-ins included in every sample clustered very tightly together, demonstrating no batch effect. Indeed, there was a very high inter-batch similarity when correlating the reference sample (0.99 ≥ *r* ≤ 1.0) or the ERCC spike-ins (0.83 ≥ *r* ≤ 1.0) across batches. A Euclidean distance-based heatmap clustering with the top 500 highly variable genes (HVG) across all samples further demonstrated a high similarity of the reference sample across batches and a tendency of AD samples to cluster separately from CTRL samples. Bulk RNA-seq data is available in the NCBI Gene Expression Omnibus (GEO) database (GSE226901).

### Quantitative immunohistochemistry

Cryostat sections adjacent to those used for LCM were subjected to multiplex fluorescent immunohistochemistry. Briefly, sections were fixed in 4% paraformaldehyde for 30 min, blocked with 10% normal donkey serum in tris-buffered saline (TBS) for 1 h at room temperature, incubated with primary antibodies in 5% normal donkey serum in TBS overnight at 4°C, washed in TBS (3×5 min), incubated with Cy3-, Cy5-, and AlexaFluor488-conjugated secondary antibodies of the appropriate species in 5% normal donkey serum in TBS for 2 h at room temperature, washed in TBS (3×5 min), counterstained with ThioS, and coverslipped with DAPI-containing mounting media (Fluoromount-G, Southern Biotech, Cat# 0100-20). Primary antibodies included a rabbit polyclonal anti-GFAP (1:1,000, Sigma-Aldrich, Cat# G9269), a mouse monoclonal anti-CD68 antibody (clone KP1, 1:50, Dako, Cat# M0814), a rabbit monoclonal anti-Aβ N-terminus (clone D54D2, 1:500, Cell Signaling Technologies, Cat# 8243S), and a mouse monoclonal anti-pTau-Ser396/404 (Dr. Peter Davies’ kind gift). Immunostained sections were scanned in a VS-120 Olympus slide scanner and area fraction for each marker (i.e., % area of cortex occupied by the immunoreactivity or ThioS staining) was estimated using the cellSens software (Olympus, Tokyo, Japan). CTRL vs. AD comparisons for each of these measures were performed with either unpaired t*-*test (normally distributed data) or Mann-Whitney U test (non-normally distributed data). All measures from AD donors were cross-correlated using Pearson’s correlation test.

### Bioinformatics analyses

#### Differential expression analyses

Raw count data was normalized using the *voom* package^30^ in R and log-transformed. Genes with low expression were filtered. The effects of age, sex, and postmortem interval (PMI) were regressed out using a linear model. Differential expression analyses were conducted using the *limma* package^31^ in R, and the models used were based on prespecified hypotheses, as follows:

1. Transcriptomic differences between AD and CTRL donors are maximum within and around Aβ plaques and NFTs:

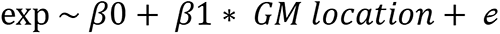 where gray matter (GM) location is a categorical variable with CTRL cortex as reference, and “plaques,” “peri-plaque,” “tangle,” and “distant” as possible values.
2. Aβ plaques and NFTs are associated with distinct transcriptomic changes and the *APOE* genotype impacts the transcriptomic changes within and around Aβ plaques and NFTs in the AD brain. We ran two separate linear regression models.
  a. A model to determine the DEGs by *APOE* genotype in each cortical location only in AD donors:

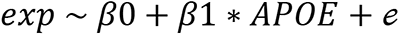 where *APOE* is a binary variable with *APOE*ε3/ε3 as reference (the single *APOE*ε3/ε4 AD donor was excluded from this analysis).
  b. A mixed-effects model to determine the effects of *APOE* genotype, cortical regions (i.e., location), and the interaction between the two:

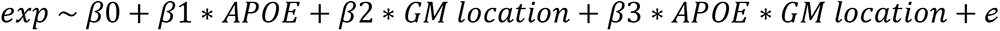

where *APOE* is a binary variable with *APOE*ε3/ε3 as reference (the single *APOE*ε3/ε4 AD donor was excluded from this analysis), *GM location* is a categorical variable with “distant” as reference, and “plaques”, “peri-plaque”, and “tangle” as possible values (CTRL donors are excluded from this analysis), and the interaction term *APOE* ∗ *GM location* represents the difference between both *APOE* genotypes at each location relative to the “distant” one. DEGs were adjusted for multiple comparisons with the Benjamini-Hochberg correction. All models were run in R (version 4.2.1).

#### Cell type assignment of differentially-expressed genes

To ascribe cell type identity to the DEGs, we took advantage of a public RNA-seq database of human brain cell subpopulations isolated by immunopanning^32^. We assigned genes to either microglia, astrocytes, neurons, oligodendrocytes, or endothelial cells, if their expression was enriched in that cell type (≥ 1.5-fold of the sum of their expression in all other cell types). This criterion rendered 3766 neuronal, 2027 microglial, 1700 astrocytic, 866 oligodendroglial, and 1050 endothelial cell-predominant genes (**Supplemental Table 1).**

#### Gene Set Enrichment Analysis (GSEA)

To investigate the possible functional changes associated with the observed transcriptomic changes, we conducted pathway enrichment analysis using GSEA^33^ against the Reactome database in the Molecular Signatures Database (MSigDB) version 3.0^34^. To eliminate redundancies across pathways, the resulting individual pathways were grouped into superpathways based on their similarity of gene composition, which was computed using the Jaccard similarity index, as below:

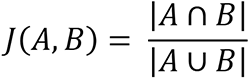

where A and B are two given Reactome pathways, |A ∩ B| is the size of the intersection between their genes, and |A ∪ B| is size of the union. The superpathways were then expert-annotated based on the overarching main function described by their constituent pathways. For each superpathway, the union of all leading-edge genes from all its Reactome pathways were compiled and subjected to GSEA to obtain the final normalized enrichment score (NES) and FDR *q* value, which were depicted in horizontal bar plots. Superpathways made of only one Reactome pathway were excluded from these plots.

Additionally, we validated previously published gene expression signatures in Aβ plaque and NFT areas by performing GSEA of those signatures against the DEGs of each of the AD cortical locations vs. CTRL cortex. The gene sets tested included: (1) the “disease-associated microglia” (DAM) signature (n=118 genes UP and 185 DOWN), which was obtained via single-cell RNA-seq of CD45+ microglia isolated from 5xFAD transgenic mice^35^; (2) the “microglial neurodegenerative phenotype” (MGnD) (n=28 UP and 68 DOWN), which was obtained via RNA-seq of FCRSL+ microglia isolated from *APP/PS1* transgenic AD mice, *SOD1*^G93A^ transgenic amyotrophic lateral sclerosis (ALS) mice, and mice with experimental autoimmune encephalomyelitis (EAE, a model of multiple sclerosis)^36^; (3) the “activated response microglia” (ARM, n=68 UP and 25 DOWN), “interferon response microglia” (IRM, n=100 UP genes), and “transiting response microglia” (TRM), which were derived via single-cell RNA-seq of Cd11b+ microglia isolated from *App^NL-G-F^* knock-in AD transgenic mice^37^; (4) the “Aβ plaque-induced genes” (PIG, n=57 UP genes) and the “oligodendrocyte-induced genes” (OLIG) modules obtained via spatial transcriptomics in the *App^NL-G-F^* knock-in AD transgenic mice^19^; (5) a microglia-*APOE* signature obtained via spectral clustering reanalysis of the ROSMAP dorsolateral prefrontal cortex bulk RNA-seq data from individuals with no neuritic plaques (CERAD NP score 0) or frequent neuritic plaques (CERAD NP score 3) across *APOE* genotypes^10^; (6) a tangle-bearing neuron signature obtained from ThioS+/NEUN+ vs. ThioS-/NEUN+ neuronal cell bodies from human AD donors^38^; (7) a pan-neurodegenerative (PAN) signature obtained from a meta-analysis of 60 bulk microarray studies encompassing more than 2,600 samples from AD, Lewy body diseases (LBD), and ALS-frontotemporal dementia (FTD) spectrum and CTRL individuals^39, 40^; (8) an AD reactive astrocyte (ADRA) signature compiled from a systematic review of postmortem immunohistochemical studies^41^; (9) a pan-injury astrocyte signature (Astro_PAN) resulting from meta-analyzing astrocyte-specific transcriptomic datasets of mouse models of acute CNS injury and chronic neurodegenerative diseases^42^; and (10) several senescence (SEN) signatures^43–45^.

#### Transcription Factor Enrichment Analysis (TFEA)

Differentially upregulated (log fold-change [FC] ≥ 1.2) and downregulated (logFC < -1.2) genes in each AD cortical region vs. CTRL cortex were input in EnrichR Chromatin Immunoprecipitation (ChIP) Enrichment Analysis (ChEA) 2022 to investigate putative transcription factors driving those gene expression changes^46–48^.

## RESULTS

### Spatial transcriptomics in human postmortem brains via LCM of A**β** plaques and NFTs

To determine whether Aβ plaques or NFTs are associated with different transcriptomic responses in the AD brain with respect to the CTRL cortex, we compared the transcriptome of laser-capture microdissected (i) ThioS+ plaques (“plaque”), (ii) the 50 μm perimeter around them (“peri-plaque”), (iii) ThioS+ NFTs plus the 50 μm perimeter around them (“tangle”), and (iv) areas devoid of dense-core ThioS+ Aβ plaques and NFTs (“distant”) from the superior temporal gyrus (STG) of n=10 AD donors, with the transcriptome of laser-captured microdissected plaque-and tangle-free cortex from n=8 CTRL donors (**Figure 1A**). Table 1 depicts the demographic, *APOE* genotype, and neuropathological characteristics of the study donors. The 50 μm perimeter was chosen because our prior stereology-based quantitative neuropathological studies in the temporal association cortex demonstrated prominent astrocyte and microglial responses to Aβ plaques and NFTs within this distance^4–7^. The STG was selected because it is an area of severe neuritic dense-core Aβ plaque deposition and NFT spreading in Braak V-VI AD donors^4–7, 49^. Quantitative measurements of Aβ-, pTau (PHF1)-, GFAP-, and CD68-immunoreactive area fraction and ThioS-stained area fraction in adjacent cryostat sections of the STG confirmed statistically significantly higher levels of Aβ plaques, NFTs, and astrocytic and microglia responses in AD vs. CTRL donors (**Figure 1B**). We observed a stronger correlation of reactive glia measures with the pTau burden than with the Aβ plaque burden (**Figure 1C**), as reported previously^4^.

**Figure 1.**
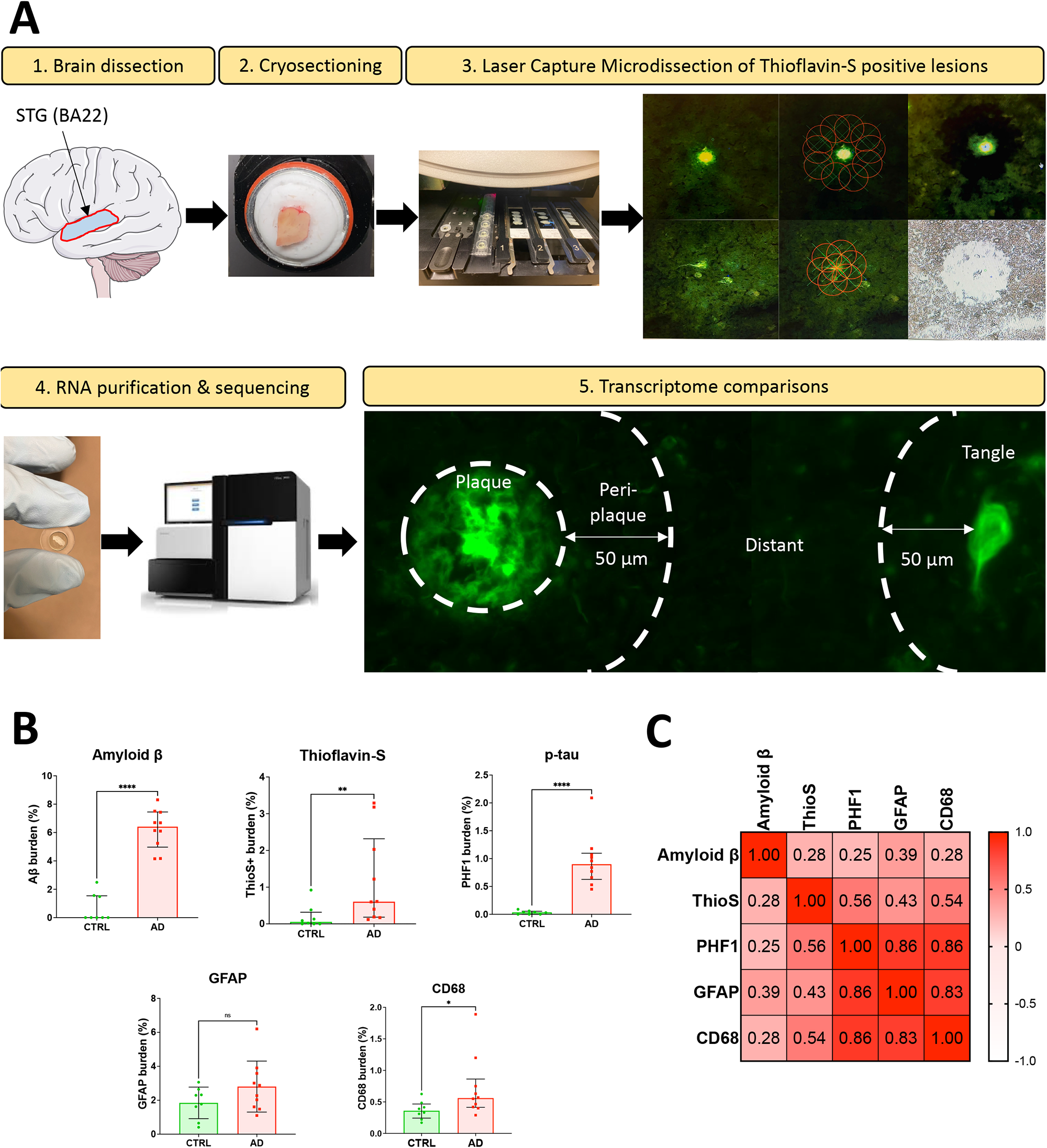
Methodology of laser capture microdissection and RNA-sequencing. (A) Cryostat sections were obtained from superior temporal gyrus (STG, BA22) and stained with Thioflavin-S (ThioS). ThioS+ Aβ plaques and NFTs were identified morphologically and laser-capture microdissected as shown; areas distant (>50 μm) from the Aβ plaques and NFTs were laser-capture microdissected as well. The transcriptomes of ThioS+ Aβ plaques, the 50 μm peri-plaque halo, ThioS+ NFTs with their 50 μm halo, and areas beyond 50 μm from the nearest ThioS+ Aβ plaque or NFT (distant) of AD donors were compared to the transcriptome of normal-appearing cortex of control individuals. The transcriptome of ThioS+ Aβ plaques was also compared with that of ThioS+ NFTs from the same AD donors. (B) Comparison of Aβ plaque, ThioS+, pTau (PHF1+), reactive astrogliosis (GFAP+) and reactive microglia (CD68+) area fractions (i.e., % cortical area occupied by immunoreactive or staining signal) in cryostat sections adjacent to those used for LCM between CTRL and AD donors (unpaired t-test or Mann-Whitney U test, * *p*<0.05, ** *p*<0.01, **** *p*<0.0001, ns = non-significant). (C) Correlation matrix heatmap illustrates the strength of correlation (Pearson’s correlation coefficients) across all quantitative measures within the AD group.

**Table 1.** Demographic, clinical, and neuropathological characteristics, and APOE genotype of study subjects.

| **Donor ID** | ***APOE* genotype** | **Age (y)** | **Sex** | **PMI (h)** | **Thal amyloid phase** | **Braak NFT stage** | **CERAD NP score** |
| --- | --- | --- | --- | --- | --- | --- | --- |
| CTRL1 | ε3/ε3 | 73 | F | 20 | 0 | 0 | None |
| CTRL2 | ε3/ε3 | 88 | M | 14 | 0 | III | None |
| CTRL3 | ε3/ε3 | 76 | F | 39 | 2 | I | Sparse |
| CTRL4 | ε3/ε4 | 77 | F | 72 | 1 | I | Sparse |
| CTRL5 | ε3/ε3 | 90+ | M | 39 | 3 | III | Sparse |
| CTRL6 | ε3/ε3 | 79 | F | 9 | 0 | II | None |
| CTRL7 | ε3/ε3 | 90+ | M | 21 | 0 | II | None |
| CTRL8 | ε2/ε3 | 87 | M | 21 | 3 | II | Sparse |
| AD1 | ε3/ε3 | 60 | F | NA | 4 | VI | Frequent |
| AD2 | ε3/ε3 | 86 | M | 20 | 5 | VI | Frequent |
| AD3 | ε3/ε3 | 85 | M | 39 | 4 | V | Moderate |
| AD4 | ε3/ε3 | 72 | F | 10 | 5 | VI | Frequent |
| AD5 | ε4/ε4 | 84 | M | NA | 5 | VI | Frequent |
| AD6 | ε4/ε4 | 71 | F | 21 | 5 | VI | Frequent |
| AD7 | ε4/ε4 | 82 | M | 10 | 4 | V | Frequent |
| AD8 | ε4/ε4 | 74 | M | 10 | 5 | V | Frequent |
| AD9 | ε4/ε4 | 75 | F | 18 | 4 | VI | Frequent |
| AD10 | $\epsilon 3/\epsilon 4$ | 85 | F | 24 | 3 | V | Moderate |

Abbreviations: F = female; M = male; NA = not available; NFT = neurofibrillary tangle; NP = neuritic plaque; PMI = postmortem interval.

### A**β** plaque microenvironment is associated with greater transcriptomic changes than NFT microenvironment

To compare and contrast the transcriptomic changes in the four locations, we performed differential expression analyses of each of the cortical locations in AD vs. CTRL cortex. The ThioS+ Aβ plaques in AD cortex had the highest number of differentially expressed genes (DEG) with 2,623 upregulated genes and 2,563 downregulated genes relative to CTRL cortex (*p*-value < 0.05) (**Figure 2AB** and **Supplemental Table 2**). Interestingly, the “peri-plaque” (that is, the 50 μm band around the edge of ThioS+ Aβ plaques) vs. CTRL cortex comparison followed with 1,681 and 1,535 unique upregulated and downregulated genes, respectively. By contrast, “tangle” and “distant” vs. CTRL cortex analyses produced similar numbers of unique upregulated (1,230 and 1,261, respectively) and downregulated (1,056 and 1,103, respectively) genes. The number of DEGs in AD vs. CTRL that were common to all AD cortical locations (631 upregulated and 494 downregulated) was about half the number of unique genes differentially expressed in each particular location, thus confirming that there are many transcriptomic changes that are specifically occurring within or in the vicinity of AD pathological hallmarks. Only the “plaque” vs. CTRL cortex comparison revealed a sizable set of differentially expressed genes after multiple comparison corrections (1,152 upregulated genes and 827 downregulated genes, adj. *p*-value < 0.05), again demonstrating that the impact of AD on cortical gene expression is strongest within the ThioS+ Aβ plaques.

### Cell type assignment of differentially-expressed genes implicates microglia in Aβ plaques and neurons and oligodendrocytes in cortical regions far from plaques and tangles

Next, we investigated the cell type distribution of the transcriptomic changes observed in each AD cortical location vs. CTRL cortex. In the Aβ plaques, most of the upregulated genes were microglial, followed by non-cell-type-specific genes (i.e., similarly expressed by multiple cell types, including microglia as well as other cells) (**Figure 2C**). In the peri-plaque areas, most upregulated genes were astroglial (excluding non-cell-type-specific genes), followed by neuronal and microglial. In NFTs, most upregulated genes were neuronal, followed by astroglial and endothelial. Finally, in areas distant from Aβ plaques and NFTs, most upregulated genes were oligodendroglial, suggesting a possible response of oligodendrocytes to axonal injury far from Aβ plaques and NFTs (**Figure 2C**). Interestingly, most downregulated genes in Aβ plaques, peri-plaque areas, NFTs as well as areas distant from ThioS+ Aβ plaques and NFTs were neuronal, pointing to significant neuronal dysfunction also in areas far from AD lesions (**Figure 2C**).

**Figure 2.**
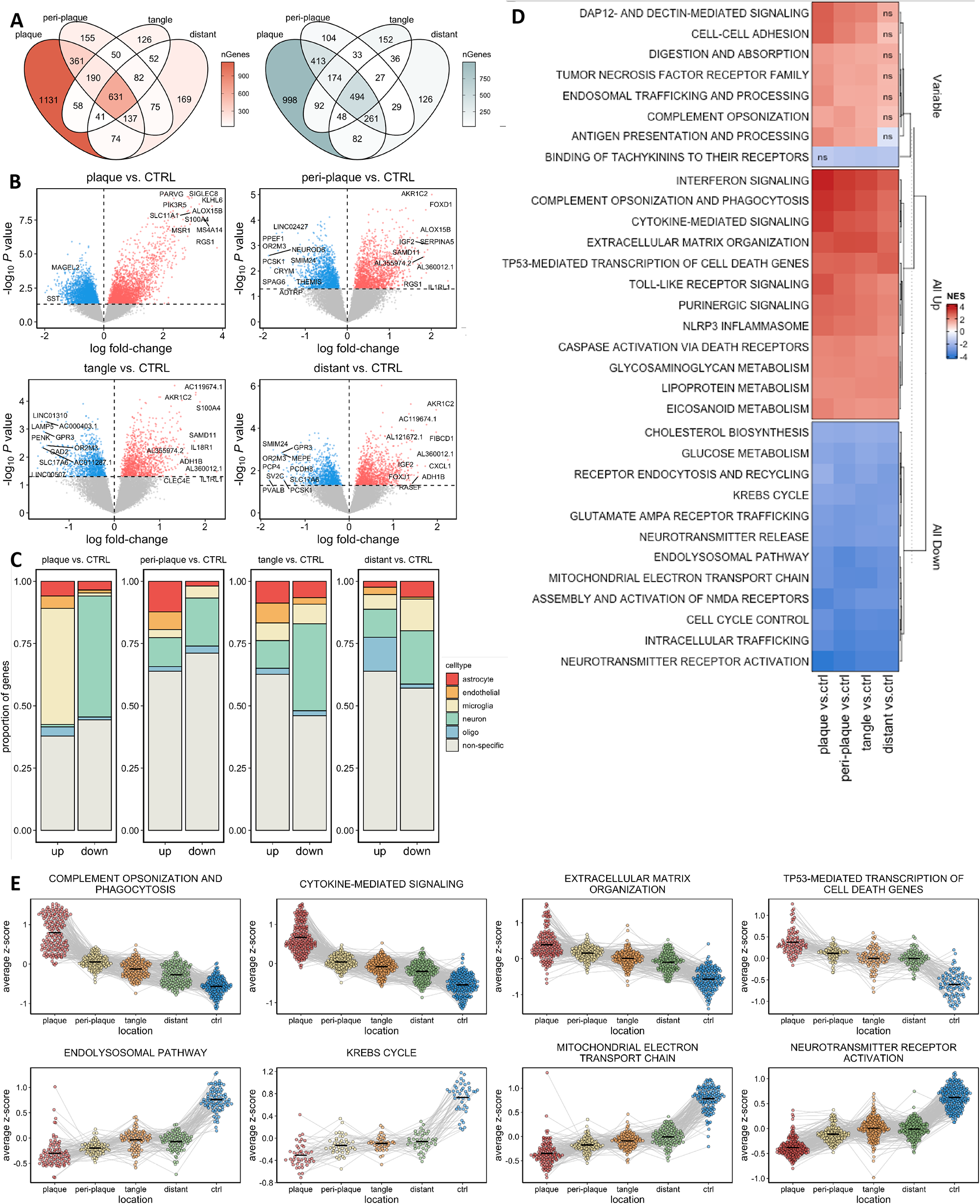
Aβ plaque microenvironment is associated with greater transcriptomic changes than NFT microenvironment. (A) Venn diagrams depict the number of differentially expressed genes (DEGs, UP in red and DOWN in blue, *p*-value < 0.05) in each AD cortical location vs. CTRL cortex. Note that the majority of transcriptomic changes occur within Thioflavin-S-positive Aβ plaque areas. (B) Volcano plots show individual top UP and DOWN DEGs in each AD cortical location vs. CTRL cortex. (C) Proportion plots showing enrichment in brain cell types of the UP and DOWN DEGs based on a public immunopanning-based human brain RNA-seq dataset (www.brainrnaseq.org). Most UP genes in Aβ plaque areas are microglial, whereas most DOWN genes across cortical locations are neuronal (gray = not cell-type specific). (D) Heatmap illustrates the functional pathway analysis of the DEGs for each AD cortical location vs. CTRL cortex. Most UP pathways are related to neuroinflammation, extracellular matrix organization, and cell death, whereas most DOWN pathways are related to neurotransmission, intracellular trafficking, and mitochondria/energy metabolism. (E) Violin plots show the expression of the leading-edge genes of relevant functional pathways connected across cortical locations. Note the gradient from CTRL cortex toward Thioflavin-S-positive Aβ plaque areas in AD cortex, either up-trending or down-trending.

Taken together, these data show that Aβ plaques trigger the bulk of transcriptomic changes occurring in the AD cortex, with significant upregulation of microglial and downregulation of neuronal genes, but also reveal the contribution of other cell types to local transcriptomic changes, notably astrocytes in peri-plaque areas and oligodendrocytes in areas distant from plaques and tangles.

### Aβ plaques are associated with greater neuroinflammation and synaptic and metabolic dysfunction than NFTs

To better understand the possible functional alterations associated with these transcriptomic changes, we conducted pathway enrichment analysis using gene set enrichment analysis (GSEA)^33^ against the Reactome database (**Figure 2D** and **Supplemental Table 3**). Among the upregulated pathways in Aβ plaques vs. CTRL cortex, those related to neuroinflammation (e.g., cytokine-mediated signaling, NLRP3 inflammasome, Toll-like receptor signaling, interferon signaling, eicosanoid metabolism) and phagocytosis (e.g., complement opsonization, antigen presentation and processing, scavenger receptors, DAP12- and Dectin-mediated signaling), cholesterol metabolism (e.g., lipoprotein metabolism, cholesterol ester synthesis), and extracellular matrix (e.g., proteoglycans and glycosaminoglycan metabolism) stood out. Additionally, TP53-regulated transcription of cell death genes, including TRAIL and death receptors, emerged as a possible mediator of plaque toxicity, together with a repression of the WNT/beta-catenin pathways and activation of NOTCH and RUNX2 and 3. By contrast, the downregulated pathways were related to neurotransmission (e.g., neurotransmitter release, neurotransmitter receptor activation, NMDA receptor assembly, AMPA receptor trafficking, depolarization), energy metabolism and mitochondrial function (e.g., Krebs cycle, electron transport chain, ATP synthesis, mitophagy, mitochondrial protein translation), intracellular trafficking (e.g., ER-to-Golgi transport, protein transport to plasma membrane, tubulin cytoskeleton), cholesterol biosynthesis, and cell cycle control (e.g., G2-M checkpoints, negative regulation of NOTCH and Sonic Hedgehog).

Of note, pathways analysis of NFTs vs. CTRL cortex DEGs only rendered a few significantly upregulated pathways, which included TP53-regulated expression of cell death genes, phagocytosis, and extracellular matrix organization, but not the prominent pro-inflammatory response described above for Aβ plaques. However, the downregulated pathways were very similar to those in the Aβ plaques. The genes within the dysregulated pathways revealed a prominent gradient with the largest changes in expression (relative to CTRL cortex) in Aβ plaques, followed by peri-plaque areas, then tangles, and then plaque/NFT-distant AD cortical areas (**Figure 2E**).

To directly compare the Aβ plaques and NFTs, we performed differential gene expression analysis between these two cortical locations in the 10 AD donors (**Supplemental Table 4**). The genes significantly higher in Aβ plaques relative to NFTs outnumbered those that were higher in the NFTs. Many of the genes that were higher in Aβ plaques were microglial (e.g., *CD84*, *CD180*, *MS4A6*, *MS4A14*, *MSR1*) (**Figure 3A**). Pathway analysis of Aβ plaques vs. NFTs showed that neuroinflammation, extracellular matrix, and cell death are enriched in Aβ plaques, whereas neurotransmission and mitochondrial electron transport chain are enriched in NFTs (**Figure 3B**). The expression levels of representative DEGs from each of these pathways are shown at the donor level in **Figure 3CD**.

**Figure 3.**
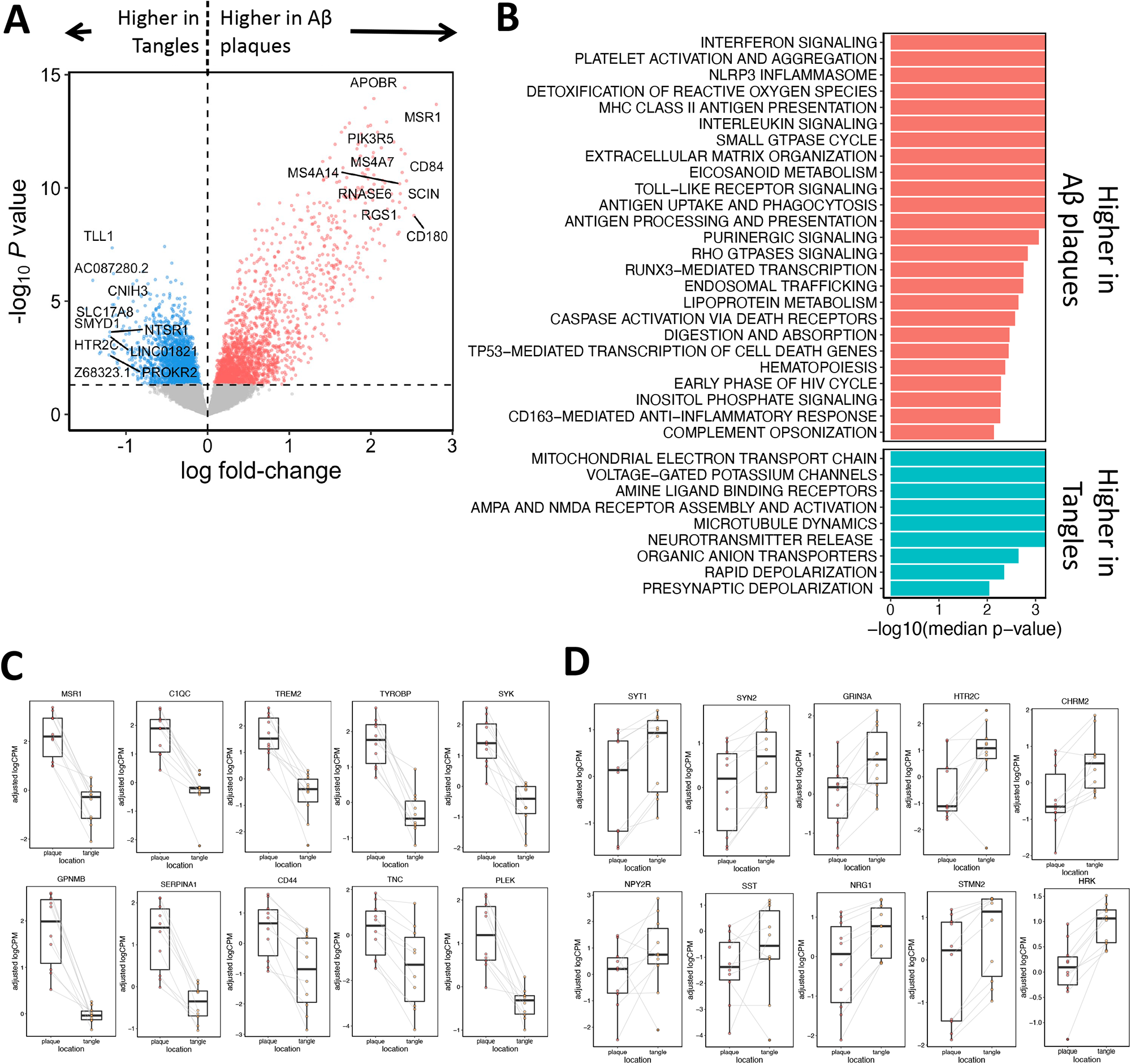
Direct comparison of Aβ plaques vs. NFTs in AD donors reveals relevant transcriptomic differences. (A) Volcano plot shows individual top DEGs in Aβ plaques vs. NFTs from AD donors. (B) Horizontal bar plot depicts the results of the functional pathway enrichment based on the Aβ plaques vs. NFTs DEGs. Note that neuroinflammation, extracellular matrix, and cell death are more enriched in Aβ plaques, whereas neurotransmission and mitochondrial electron transport chain are more enriched in NFTs. (C-D) Box plots showing level of expression of representative genes significantly upregulated in Aβ plaques vs. NFTs (C) or NFTs vs. Aβ plaques (D).

To identify putative transcription factors driving these transcriptomic responses, we performed transcription enrichment factor analysis (TFEA) by interrogating the recently updated database of ChIP experiments ChEA 2022 with the DEGs for each cortical AD location vs. CTRL cortex. These analyses suggested a remarkable local diversity of transcription factors implicated in the aforementioned transcriptomic changes. In particular, the microglial SPI1 (a.k.a. PU.1) was the main enriched transcription factor associated with upregulated genes (i.e., transcriptional activator) in Aβ plaque areas, the NFκB subunit RELB in peri-plaque areas, and ELK3 in NFT areas, whereas NEUROD2 and TP53 (a.k.a. p53) were the most enriched in areas distant from plaques and NFTs. On the other hand, SUZ12 and REST emerged as the main transcription factors associated with downregulated genes (i.e., transcriptional repressors), particularly in Aβ plaque and distant areas, whereas the retinoic acid receptor-β (RARB) and the proto-oncogene MYC were the top repressors in NFTs (**Figure 4** and **Supplemental Table 5**).

**Figure 4.**
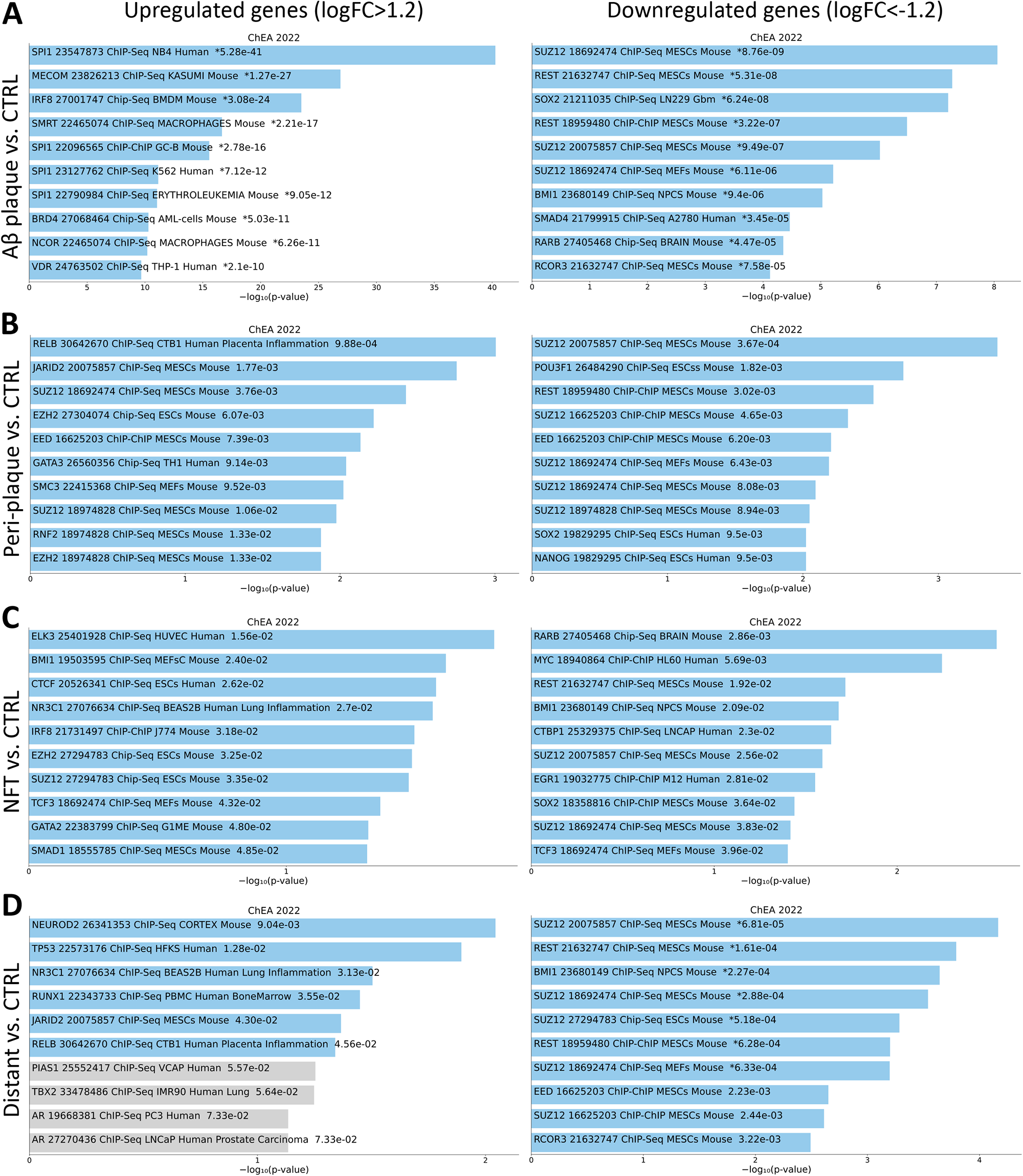
Distinct transcription factors drive local transcriptomic changes in AD cortex. Results of the transcription factor enrichment analysis of differentially upregulated (logFC ≥ 1.2) and downregulated (logFC < -1.2) genes in each AD cortical location vs. CTRL cortex based on Enrichr ChEA 2022 database (https://maayanlab.cloud/Enrichr/). Bars are sorted by *p*-value ranking. Note that SPI1 (a.k.a. PU.1) is the top transcription factor associated with upregulated genes in Aβ plaques (A), RELB in peri-plaque areas (B), ELK3 in NFTs (C), and NEUROD2 and TP53 in areas distant from Aβ plaques and NFTs (D), whereas SUZ12 is the top transcriptional repressor in Aβ plaques and peri-plaque and distant areas followed by REST in Aβ plaques and distant areas, and RARB and MYC are the top repressors in NFTs.

### The *APOE*ε4 allele exacerbates transcriptomic changes in Aβ plaques and NFTs

Next, we investigated the effects of the *APOE* genotype on transcriptomic changes across the four cortical locations in AD donors. First, we performed differential gene expression analysis between *APOE*ε3/ε3 (n=4) vs. *APOE*ε4/ε4 (n=5) AD donors for each cortical location (see Methods). We found a total of 324 upregulated and 367 downregulated genes in *APOE*ε4/ε4 vs. *APOE*ε3/ε3 AD donors common to all locations (adj. *p*-value <0.05). Additionally, we found 244/329, 165/213, 56/70, and 341/287 up/down genes in *APOE*ε4/ε4 vs. *APOE*ε3/ε3 AD donors that were unique to Aβ plaques, peri-plaque, NFTs, and distant locations, respectively (**Figure 5AB** and **Supplemental Table 6**). Pathway analysis on the *APOE*ε4/ε4 vs. *APOE*ε3/ε3 DEGs in Aβ plaques demonstrated an upregulation of innate immune and pro-inflammatory pathways (cytokine-mediated signaling, interferon signaling, NFkB activation, NLRP3 inflammasome, complement opsonization and phagocytosis), cell death pathways (caspase-activation via death receptors, TP53-mediated transcription of cell death genes, chaperone-mediated autophagy, and RIPK1-mediated necroptosis), protein translation, extracellular matrix organization, and cholesterol metabolism, and a downregulation of neurotransmission (**Figure 5C** and **Supplemental Table 7**). Thus, relative to the *APOE*ε3 allele, the *APOE*ε4 allele exacerbated the transcriptomic changes of Aβ plaques vs. normal cortex described above. Similarly, compared to *APOE*ε3 homozygotes, *APOE*ε4 homozygotes had an upregulation of pro-inflammatory (particularly interferon and interleukin-mediated signaling, but less prominent compared to plaques), phagocytosis, cell death, protein translation, extracellular matrix, and cholesterol metabolism, and a downregulation of neurotransmission and synaptic pathways in NFT areas (**Figure 5C**).

**Figure 5.**
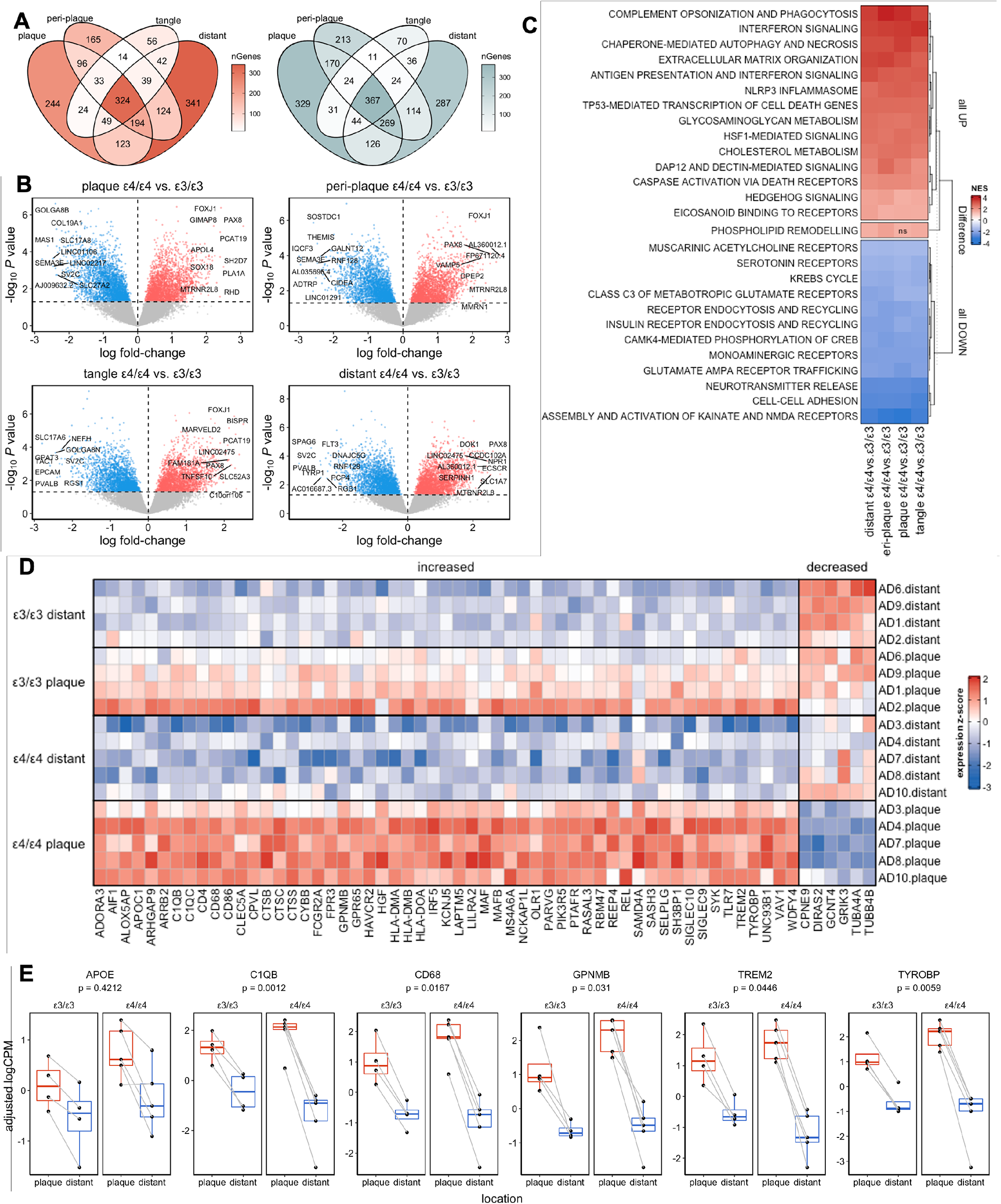
The *APOE*ε4 allele exacerbates transcriptomic changes in both Aβ plaques and NFTs. (A) Venn diagrams depict the number of DEGs (UP in red and DOWN in blue, *p*-value < 0.05) in AD *APOE*ε4 vs. *APOE*ε3 homozygotes for each cortical location. (B) Volcano plots show individual top UP and DOWN DEGs in AD *APOE*ε4 vs. *APOE*ε3 homozygotes for each cortical location. (C) Heatmap illustrates the functional pathway enrichment of the DEGs in AD *APOE*ε4 vs. *APOE*ε3 homozygotes for each cortical location. Note that most UP pathways in AD *APOE*ε4 vs. *APOE*ε3 homozygotes are related to neuroinflammation, extracellular matrix organization, and cell death, whereas most DOWN pathways are related to neurotransmission, intracellular trafficking, and mitochondria/energy metabolism across and that this is the case across cortical locations. (D) Heatmap depicts 61 DEGs (55 UP and 6 DOWN) that were significantly different in AD *APOE*ε4 vs. *APOE*ε3 homozygotes in Aβ plaque vs. distant locations. Note that most *APOE*ε4-associated UP genes were microglial and that the *APOE*-*TREM2*-*TYROBP* axis emerged in Aβ plaques. (E) Box plots show the AD donor-level expression of representative *APOE*ε4-associated genes in plaque vs. distant locations by *APOE* genotype. Note that all these genes were upregulated in Aβ plaque vs. distant locations and, except *APOE* itself, also further upregulated in AD *APOE*ε4 vs. *APOE*ε3 homozygotes (*p*-value denotes the level of statistical significance of the *APOE* × cortical location interaction term).

Next, to further investigate the impact of *APOE* genotype on relative gene expression across spatial locations within donors, we applied mixed-effects models with *APOE* genotype (*APOE*ε3/ε3 as reference), cortical location (“distant” as reference), and interaction terms between *APOE* and cortical location (see Methods). We identified 55 genes that were significantly upregulated within the Aβ plaques relative to areas distant from both plaques and NFTs and further upregulated in *APOE*ε4/ε4 vs. *APOE*ε3/ε3 donors, whereas only 6 genes were significantly downregulated within the Aβ plaques vs. distant areas and further downregulated in *APOE*ε4/ε4 vs. *APOE*ε3/ε3 donors (**Figure 5D**). Notably, most of these 55 upregulated genes were microglial, including two AD risk genes (*MS4A6A* and *TREM2*), inflammatory (*ALOX5AP*, *GPNMB*, *HAVCR2*, *IRF5*, *REL*, *TLR7*, *UNC93B1*) and phagocytosis genes (cell motility [*AIF1*, *NCKAP1L*, *PARVG*, *SH3BP1*, *VAV1*], cell adhesion [*ARHGAP9*, *SELPLG*], antigen presentation [*CD4*, *CD86*, *FCGR2A*, *HLA-DMA*, *HLA-DMB*, *HLA-DOA*, *LILRA2*], opsonization [*C1QB*, *C1QC*], phagocytosis receptors and adaptors [*CLEC5A*, *SIGLEC9*, *SIGLEC10*, *SYK*, *TREM2*, *TYROBP*], respiration burst [*CYBB*], lysosomal [*CD68*, *CPVL*, *CTSB*, *CTSC*, *CTSS*, *LAPTM*]), and a few lipid metabolism genes (e.g., *APOC1* and *OLR1*) (**Figure 5E**). By contrast, we found no significant interaction between *APOE* genotype and peri-plaque and NFT locations, suggesting that the *APOE*ε4 allele has the greatest impact on the transcriptome of Aβ plaque-associated microglia.

### AD glial and neuronal transcriptomic signatures are variably enriched in Aβ plaques vs. NFTs

Given that microglial genes and pathways were over-represented in Aβ plaques vs. CTRL cortex, we investigated whether several microglial transcriptomic signatures previously reported in AD mouse models and bulk brain from individuals with AD are present in Aβ plaques from human AD patients. Specifically, we tested the disease-associated microglia (DAM)^35^ signature, the microglial neurodegenerative phenotype (MGnD)^36^, the activating response microglia (ARM)^37^, the interferon response microglia (IRM)^37^, the transiting response microglia (TRM)^37^, the “plaque-induced genes” (PIGs) module^19^, and the microglia-*APOE* signature^10^ via GSEA of these gene sets against the set of DEGs resulting from the Aβ plaques and NFTs vs. normal-appearing CTRL cortex comparisons (**Figure 6A** and **Supplemental Table 8**). Of all these microglial signatures, the microglia-*APOE*, PIGs, MGnD UP and DOWN, IRM, ARM UP and DOWN, and DAM UP were significantly enriched in Aβ plaques vs. CTRL cortex, whereas only MGnD UP was significantly enriched in NFTs vs. CTRL cortex.

**Figure 6.**
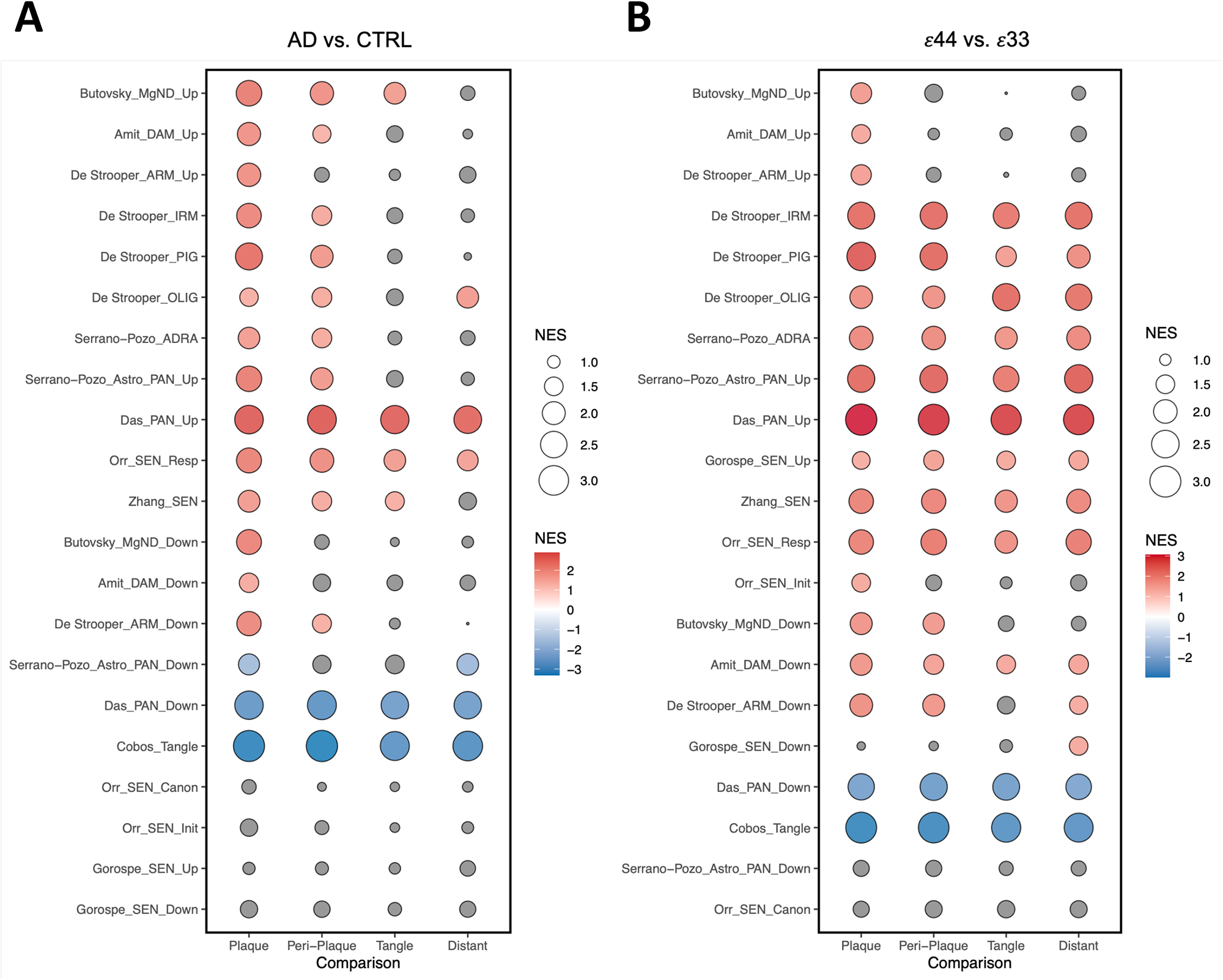
Published AD glial and neuronal signatures are variably enriched in Aβ plaques vs. NFTs. Bubble charts depict the normalized enrichment score (NES) of the gene set enrichment analysis (GSEA) of each of the published signatures (left names) against the DEGs between (A) each of the AD cortical locations and CTRL cortex, and (B) AD *APOE*ε4 vs. *APOE*ε3 homozygotes in each cortical location. Red indicates statistically significant enrichment; blue statistically significant anti-enrichment; and gray non-statistically significant result.

Additionally, we investigated other cell-type specific signatures reported in the literature. An AD reactive astrocyte (ADRA) signature obtained from a systematic review of the neuropathological literature^41^ and a pan-injury astrocyte upregulated signature (Astro_PAN_Up) derived from a meta-analysis of astrocyte transcriptomic studies of mouse models of acute CNS injury and chronic neurodegenerative diseases^42^ were enriched in Aβ plaques and peri-plaque areas, whereas the Astro_PAN_Down was down-regulated in Aβ plaques and distant areas. An oligodendrocyte (OLIG) module found in a spatial transcriptomic study in *App*^NL-G-F^ knock-in AD transgenic mice^19^ was enriched in Aβ plaques, peri-plaque, and distant areas but not in NFTs. A pan-neurodegenerative upregulated (PAN_Up) signature obtained from a meta-analysis of more than 2,600 samples from AD, LBD, and ALS-FTD and CTRL individuals^39, 40^ and a curated senescence response signature (SEN-Resp) containing many components of the senescence-associated secretory phenotype (SASP) and reported to be enriched in AD excitatory neocortical neurons^44^ were enriched in all cortical locations, whereas a senescence gene set highly conserved across 50 human tissues^45^ was enriched in all cortical locations except for areas distant to Aβ plaques and NFTs. Surprisingly, a tangle-specific signature (Tangle) obtained from single-cell RNA-seq of ThioS+ NFT-bearing neurons vs. NFT-free neurons^38^ was significantly downregulated across cortical locations and most within Aβ plaques. The pan-neurodegenerative downregulated (PAN_Down) signature^39, 40^ was also significantly downregulated across cortical locations. Lastly, an *in vitro* senescence signature (SEN_Up and SEN_Down) obtained from four different senescence cell models^43^ and the canonical (SEN_Canon) and initiating (SEN_Init) senescence signatures reported to be enriched in AD excitatory neocortical neurons^44^ were not significantly enriched in any cortical location.

In the *APOE*ε4/ε4 vs. *APOE*ε3/ε3 contrast, these previously identified gene expression signatures had higher normalized enrichment scores and were statistically significant across more cortical locations (**Figure 6B** and **Supplemental Table 8**), suggesting that the *APOE*ε4 allele exacerbates these altered functional pathways.

## DISCUSSION

The unique design of this LCM/RNA-seq study enabled us to dissect the transcriptomic changes associated with Aβ plaques vs. NFTs in the AD cortex and any *APOE*-linked differential responses to these two types of lesions. We found that Aβ plaques are a major contributor to the majority of the transcriptomic changes in the AD cortex as they are associated with a higher number of DEGs relative to the normal-appearing CTRL cortex than NFTs. Aβ plaque-associated transcriptomic changes consisted of an upregulation of predominantly microglial pro-inflammatory and phagocytosis genes, and a down-regulation of predominantly neuronal neurotransmission, energy metabolism, and mitochondrial function genes. This signature is remarkably similar to the pan-neurodegenerative signature that we identified in a meta-analysis of bulk transcriptomic studies from more than 2,600 postmortem samples from AD, LBD, and ALS/FTD and CTRL individuals^39, 40^, reinforcing the idea that ThioS+ Aβ plaques leave a neurodegenerative footprint in the cortical neuropil. NFTs-associated transcriptomic changes also consisted of an upregulation of microglial phagocytic genes—although neuroinflammation was not significant in the functional enrichment analysis—and a downregulation of neuronal genes involved in neurotransmission, energy metabolism, and mitochondrial function. Notably, NFTs-associated changes were less pronounced than those in Aβ plaques as indicated by fewer DEGs and by the within-AD-donor comparison showing greater upregulation of pro-inflammatory pathways and greater downregulation of neurotransmission and mitochondrial/energy metabolism pathways in Aβ plaques relative to NFTs.

We investigated the contributions of various brain cell types to the observed upregulated and downregulated genes in each AD cortical location vs. CTRL cortex. While microglia was the predominant cell type driving the upregulated transcriptional responses within Aβ plaques, astrocytes exhibited a more prominent role in the peri-plaque areas. Importantly, these findings are consistent with classic neuropathological reports describing that the cell bodies of reactive microglia are often located within dense-core Aβ plaques, whereas the cell bodies of reactive astrocytes (i.e., where most mRNA is) are usually located in the peri-plaque area and their processes surround and penetrate the plaques^5, 50^. Also, in agreement with those observations, the analyses suggest that the pro-inflammatory and anti-phagocytic SPI1/PU.1^51, 52^ was the top transcription factor driving upregulated genes in Aβ plaques, whereas the catalytic subunit of NFkB RELB, which is involved in reactive astrogliosis^53^, was the top transcription factor driving upregulated genes in peri-plaque areas. On the other hand, consistent with widespread neuron loss in advanced AD, neurons were the main cell type driving the downregulated transcriptional programs across all locations in the AD cortex, particularly within ThioS+ dense-core Aβ plaques, which are known to cause focal neuron loss^54^. Interestingly, SUZ12 and REST emerged as the main transcriptional repressors in Aβ plaque, peri-plaque, and distant areas in our transcription factor enrichment analysis. REST has been attributed a neuroprotective role through both repressing cell death genes and promoting the expression of anti-oxidant genes, and was reported to be depleted in AD brains; in addition, both SUZ12 and REST have been linked with a dysregulation of the neuronal plasticity protein network in AD in a CSF proteomics study^55^. In NFTs, RARB and MYC were the top transcriptional repressors. Expression of cell cycle markers by NFT-bearing neurons is a long-known phenomenon^56, 57^ and overexpression of the proto-oncogen MYC by excitatory neurons induces cell-cycle re-entry which results in neuronal death^58^. Moreover, pharmacological inhibition of MYC in neurons from a tauopathy mouse model has been shown to be neuroprotective^59^. Intriguingly, most of cell-type specific genes upregulated in plaque/NFT-distant areas vs. CTRL cortex were oligodendroglial, suggesting a response of oligodendrocytes to ongoing axonal damage. The role of oligodendrocytes and axon myelination in AD has been also highlighted by other recent transcriptomic studies in human AD brains and transgenic AD _mice_^12,18,19,60^.

Our dataset allowed us to investigate the mechanisms of neuronal death from a transcriptomic perspective. The extrinsic pathway of apoptosis, specifically TP53-mediated transcription of cell death genes (including death receptors and ligands) and caspase-activation via death receptors emerged as the main cell death pathways in all AD cortical locations vs. CTRL cortex according to our pathway analysis. Moreover, the tumor suppressor factor TP53 (a.k.a. p53) was one of the top transcription factors in plaque/NFT-distant cortical areas based on our TFEA. Of note, p53 has been described as a driver of neurodegeneration in *C9orf72*-linked ALS^61^ and also as a mediator of neuroprotective responses in tauopathy via regulation of synaptic genes^62^. MYC and p53 were among the top nodes emerging from a protein-protein interaction network of 196 markers of reactive astrocytes resulting from a systematic review of the AD neuropathological literature^41^. TP53 can also induce senescence, but of the six senescence signatures tested, only the senescence-associated secretory phenotype (SASP) was significantly enriched in AD vs. CTRL cortex^44, 45^.

Lastly, we investigated *APOE*-liked differences in transcriptomic responses to AD neuropathological hallmarks. We observed that, relative to *APOE*ε3, the *APOE*ε4 allele augments the transcriptomic changes that occur within Aβ plaques—both the upregulation of pro-inflammatory and phagocytic microglial genes and the downregulation of synaptic neuronal genes. Although our previous unbiased, stereology-based, quantitative neuropathological studies failed to detect an increase in Aβ plaque-associated IBA1+ or CD68+ microglia in *APOE*ε4 carriers vs. non-carriers^4, 5, 7^, these findings are in agreement with our recent analysis of public bulk RNA-seq datasets showing a pro-inflammatory and phagocytic microglial signature in *APOE*ε4 carriers relative to *APOE*ε3/ε3 individuals without neuritic Aβ plaques (CERAD NP score 0), which is well-preserved across these genotypes in individuals with frequent neuritic Aβ plaques (CERAD NP score 3)^10^. Indeed, we found a significant overlap (n=20) between the genes of this microglia-*APOE* signature (n=172) and the genes that are upregulated in Aβ plaques relative to distant areas and further upregulated in *APOE*ε4 homozygotes compared to *APOE*ε3 homozygotes (n=55). Significantly, among these common 20 genes were *TYROBP* and *TREM2*, thus confirming the existence of an *APOE*-*TREM2-TYROBP* axis in Aβ plaque-associated microglia in the human AD brain^10^ as described in mouse models of cerebral β-amyloidosis^35, 36^. In sharp contrast, we found a very low number of unique DEGs in *APOE*ε4 vs. *APOE*ε3 homozygotes in NFT areas and no significant interaction between *APOE* genotype and NFT location in the mixed-effect model, indicating that the differential impact of the *APOE*ε4 vs. *APOE*ε3 alleles on the transcriptome is less robust in NFTs than in Aβ plaques. This was unexpected, considering that *APOE*ε4 has been associated with tau-induced neurodegeneration in tauopathy mice via its effects on microglia gene expression^63–65^. Lastly, consistent with the microglia-*APOE* signature we identified in individuals without neuritic Aβ plaques, we also found a significant difference in gene expression in areas distant from Aβ plaques and NFTs in *APOE*ε4 vs. *APOE*ε3 homozygotes, suggesting that the *APOE*ε4 allele has a broad effect on the cortical transcriptome.

In summary, using LCM followed by RNA-seq in ThioS-stained postmortem brain sections from AD and CTRL donors, we found a gradient of transcriptomic changes from Aβ plaques to peri-plaque areas to NFTs to areas distant to plaques and NFTs. These changes consisted of an upregulation of predominantly microglial pro-inflammatory and phagocytic genes and a downregulation of predominantly neuronal synaptic, mitochondrial, and energy metabolism genes. The *APOE*ε4 allele exacerbated the microglial changes within Aβ plaques towards a pro-inflammatory and phagocytic phenotype. While spatial transcriptomics methods continue to improve toward reaching genome-wide coverage and single-cell resolution^18, 19, 66, 67^, our RNA-seq study on laser-capture microdissected samples from human AD and CTRL brains is a valuable resource to investigate the transcriptomic responses of various cell types to Aβ plaques and NFTs and the *APOE*ε4 allele effects on such responses.

## ABBREVIATIONS

Aβ: amyloid-β
AD: Alzheimer’s disease
ALS: amyotrophic lateral sclerosis
*APOE*: apolipoprotein E
ARM: activated-response microglia
BA22: Brodmann’s area 22
CTRL: control
DAM: disease-associated microglia
DEGs: differentially expressed genes
FC: fold-change
FTD: frontotemporal dementia
GSEA: gene set enrichment analysis
IRM: interferon-response microglia
LBD: Lewy body diseases
LCM: laser-capture microdissection
MGnD: microglial neurodegenerative phenotype
NFTs: neurofibrillary tangles
NP: neuritic plaque
PIGs: plaque-induced genes
pTau: phospho-tau
RIN: RNA Integrity Number
RNA-seq: RNA-sequencing
SEN: senescence
snRNA-seq: single-nucleus RNA-sequencing
STG: superior temporal gyrus
TBS: tris-buffered saline
TFEA: transcription factor enrichment analysis
ThioS: Thioflavin-S

## ACKNOWLEDGMENTS / CONFLICTS / FUNDING SOURCES / CONSENT STATEMENT

### Acknowledgements

We want to thank the patients and families involved in research at the Massachusetts Alzheimer’s Disease Research Center. We also acknowledge Patrick Dooley and Tessa Connors from the MADRC brain bank for their help providing the samples.

### Consent statement

All donors or their next-of-kin provided written informed consent for the brain donation and the present study was approved under the MGH Institutional Review Board.

## SUPPLEMENTAL MATERIAL

**Supplemental Table 1.- List of genes predominantly expressed in each cell type used for cell-type assignment**.

**Supplemental Table 2.- Differentially expressed genes in each cortical location from AD donors vs. CTRL cortex**.

**Supplemental Table 3.- Functional enrichment analysis of the differentially expressed genes in each cortical location from AD donors vs. CTRL cortex**.

**Supplemental Table 4.- Differentially expressed genes and functional enrichment analysis in Aβ plaques vs. NFTs from AD donors**.

**Supplemental Table 5. Transcription Factor Enrichment Analysis results**.

**Supplemental Table 6.- Differentially expressed genes in *APOE*ε4/ε4 vs. *APOE*ε3/ε3 AD donors in each cortical location**.

**Supplemental Table 7.- Functional enrichment analysis of differentially expressed genes in *APOE*ε4/ε4 vs. *APOE*ε3/ε3 AD donors in each cortical location**.

**Supplemental Table 8.- Results of the Gene Set Enrichment Analysis (GSEA) of relevant published signatures**.

